# Time-dependent toxicity of tire particles on soil nematodes

**DOI:** 10.1101/2021.06.29.450331

**Authors:** Shin Woong Kim, Eva F. Leifheit, Stefanie Maaß, Matthias C. Rillig

**Affiliations:** Institute of Biology, Freie Universität Berlin, 14195 Berlin, Germany; Berlin-Brandenburg Institute of Advanced Biodiversity Research. 14195 Berlin, Germany; Institute of Biochemistry and Biology, Universität Potsdam, 14476 Potsdam, Germany

**Keywords:** *Caenorhabditis elegans*, Exposure time, Lifetime, Microplastics, Soil incubation

## Abstract

Tire-wear particles (TWPs) are being released into the environment by wearing down during car driving, and are considered an important microplastic pollution source. The chemical additive leaching from these polymer-based materials and its potential effects are likely temporally dynamic, since larger amounts of potentially toxic compounds can gradually increase with contact time of plastic particles with surrounding media. In the present study, we conducted soil toxicity tests using the soil nematode *Caenorhabditis elegans* with different soil pre-incubation (30 and 75 days) and exposure (short-term exposure, 2 days; lifetime exposure, 10 days) times. Soil pre-incubation increased toxicity of TWPs, and the effective concentrations after the pre-incubation were much lower than environmentally relevant concentrations. The lifetime of *C. elegans* was reduced faster in the TWP treatment groups, and the effective concentration for lifetime exposure tests were 100- to 1,000-fold lower than those of short-term exposure tests. Water-extractable metal concentrations (Cr, Cu, Ni, Pb, and Zn) in the TWP-soils showed no correlation with nominal TWP concentrations or pre-incubation times, and the incorporated metals in the TWPs may be not the main reason of toxicity in this study. Our results show that toxic effects of TWPs can be time-dependent, both in terms of the microplastic particles themselves and their interactions in the soil matrix, but also because of susceptibility of target organisms depending on developmental stage. It is vital that future work consider these aspects, since otherwise effects of microplastics and TWPs could be underestimated.

**Graphical Abstract:** 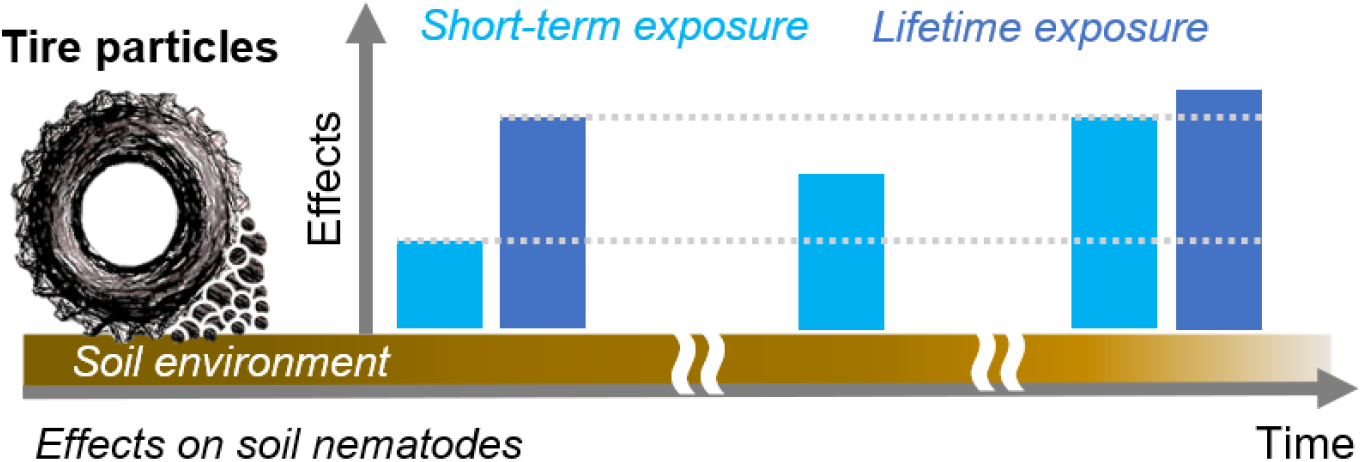

## INTRODUCTION

Since the first car tire was produced in 1895, their global production has risen due to a steady increase of the motor vehicles market (Baensch-Baltruschat et al., 2020; Halle et al., 2020). The first identification of tire rubbers in road dust was reported not before 1966 (Thompson et al., 1966), and they have been considered as a vehicle-derived pollutant (Bogdan and Albrechcinski, 1981). The modern tire consists of natural or synthetic rubbers, fillers, and resins, and these polymer-based materials can be released into the environment by wearing down with heat and friction during driving (Baensch-Baltruschat et al., 2020). Currently, tire particles are considered as microplastic due to their polymer structure, solid state, and particle sizes (Hartmann et al., 2019), and the contribution of tire particles to total microplastic emission reaches 30–50% (Baensch-Baltruschat et al., 2020). About 1.3 million tons of tire particles are generated annually in Europe (Wagner et al., 2018), and the global emission is estimated to be more than 3.3 megatons (Kole et al., 2017).

The potential toxicity of tire particles has already been highlighted in 1974 (Cardina, 1974; Dannis, 1974), and subsequent studies focused on chemical leaching of tire particles in the aquatic environment (Hartwell et al., 2000; Gualtieri et al., 2005). This is appropriate since tire particles contain a variety of leachable additives such as silica, CaCO_3_ (as fillers), polycyclic aromatic hydrocarbon (PAH, as a softener), metals, and several antioxidants (Baensch-Baltruschat et al., 2020). Leachate studies have investigated various leaching processes and test species, and delivered variable results as a function of tire composition and leachate generation times (Halle et al., 2020; Wagner et al., 2018; Wik et al., 2009). Recently, research moved from leachate-induced toxicity to particulate-induced toxicity because the ingestion and gut retention of tire particles have emerged as a crucial aspect (Halle et al., 2020). Tire particles induce toxicity on freshwater crustaceans (Khan et al., 2019; Redondo-Hasselerharm et al., 2018) and terrestrial organisms such as earthworms and plants (Leifheit et al., 2021; Pochron et al., 2017; 2018). In some studies, it seems that tire particles have high toxicity already at low concentration, but there is still a very limited number of studies, especially based on long-term exposure periods (Halle et al., 2020).

In microplastic research, many previous studies have emphasized particle-induced effects. For instance, microplastics can physically cause intestinal damage (Chae et al., 2018), and the movement of springtails can be inhibited by microplastic particles in soil pores (Kim and An, 2019). Currently, it is assumed that the chemical additives can contribute strongly to microplastic toxicity, especially when leaching out during the degradation process of particles but also already in the ‘inedible’ size ranges (Hahladakis et al., 2018; Kim et al., 2020; Zimmermann et al., 2020). The leaching of additives from polymer-based materials is highly related to both chemical equilibria and diffusion kinetics, and the leaching rate generally depends on time (Koelmans et al., 2016; Kwon et al., 2017). This means that there will be a time-delayed release of chemical additives, and larger amounts of potentially toxic compounds can be released with increasing contact time of plastic particles with the surrounding media (Rillig et al., 2021).

In the present study, we conducted soil toxicity tests using tire particles (TWPs). We prepared TWPs by grinding a used car tire, and directly mixed them with test soil. The nematode *Caenorhabditis elegans* was chosen as a model organism, and treated with TWP-soil mixtures for short-term (2 days) and lifetime (10 days) exposure periods. To evaluate the potential effects of soil pre-incubation times, we adopted three different pre-incubation times including no pre-incubation, 30 days, and 75 days (Figure 1). We analyzed water-extractable metals in the soils with different TWP concentrations and soil pre-incubation times, and evaluated time-dependent changes on TWP toxicity in the soil environment.

**Figure 1.**
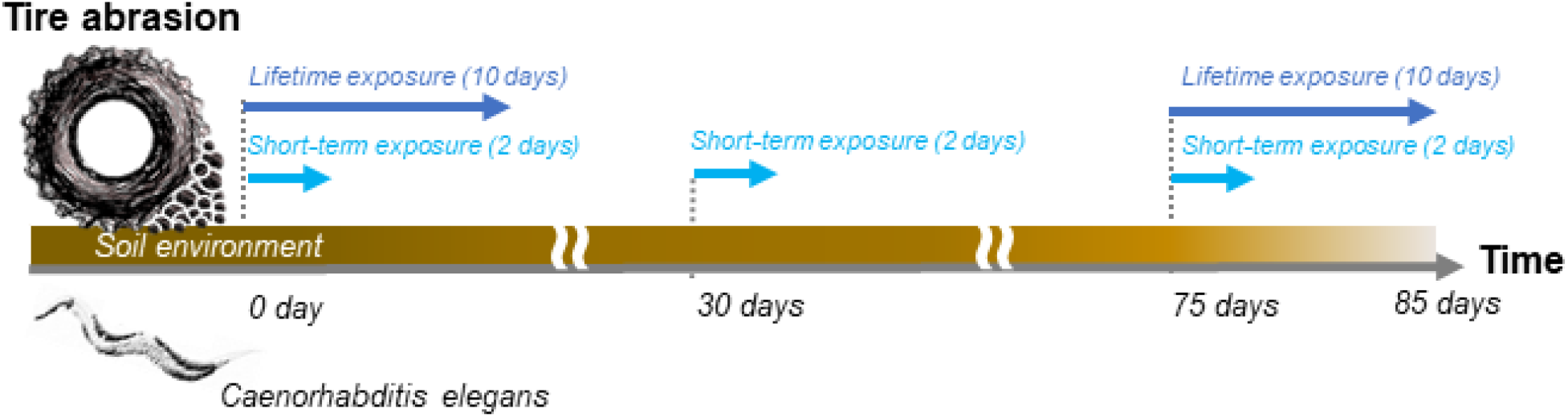
The diagram of soil toxicity tests in this study. The TWP toxicity tests were conducted using nematode *Caenorhabditis elegans* with two different exposure times (2 days and 10 days) and three different pre-incubation times (no pre-incubation, 30 days, and 75 days).

## MATERIALS AND METHODS

### Preparation and Characterization of TWPs

We prepared TWPs from a used car tire (Goodyear M+S 195/60R15 88H) using a portable belt grinding machine (Bosch PBS 75 AE). The average size of TWPs was 125 μm within a range of 34–265 μm, and heavy metal contents in TWPs were measured with an aqua regia digestion (Cr 175±7 μg g^-1^, Pb 357±29 μg g^-1^, Zn 5089±40 μg g^-1^, Ni 95±3 μg g^-1^ and Cu 453±10 μg g^-1^) (ICP-OES Perkin Elmer Optima 2100DV) (Leifheit et al., 2021). For the spectroscopic characterization of TWPs, we used a spectrophotometer (Jasco, model FT/IR-4100, ATR mode; 4000 to 600 cm^-1^, resolution of 4 cm^-1^) (Figure S1).

### Test Soil and Species

We collected test soil from a grassland site of the Institute of Biology of Freie Universität, Berlin, Germany (52.45676N, 13.30240E) on January 20, 2020. The soil was passed through a 2 mm-sieve, and dried at 60°C for 24 h. The texture of test soil was sand (sand 93.3%, silt 5.0%, and clay 1.7%), water holding capacity (WHC) was 0.34±0.10 mL g^-1^ and pH was 6.7±0.2 (Kim et al., 2021). The test soil was autoclaved twice at 121°C for 15 min to reduce microbial effects during soil toxicity tests. In order to prepare TWP-soils, 500 mg of TWPs was first mixed with 49.5 g of dry test soil (10,000 mg kg^-1^), and then this initial mixture was sequentially diluted with the same test soil. Final test concentrations were determined as 1, 10, 100, 1,000, and 10,000 mg kg^-1^, and each TWP-soil mixture was stored at room temperature.

We obtained the nematode *C. elegans* (wild type, Bristol strain N2) from the Berlin Institute for Medical Systems Biology at the Max Delbrück Center for Molecular Medicine (Berlin, Germany). The nematodes were maintained on nematode growth medium (NGM; NaCl 3 g L^-1^, peptone 2.5 g L^-1^, agar 17 g L^-1^, 1 M potassium phosphate 25 mL L^-1^, 1 M CaCl_2_·2H_2_O 1 mL L^-1^, 1 M MgSO_4_·7H_2_O 1 mL L^-1^, cholesterol 1 mL L^-1^) at 20±2°C in the dark with *Escherichia coli* (strain OP50) as a food source (Brenner et al., 1974). In order to synchronize the developmental stage, 3 day-old adults were collected, and they were treated with 3 mL of Clorox solution (1 N NaOH:5% NaOCl, 1:1) for 20 min. The suspension containing embryos was centrifuged at 5,000 rpm for 2 min, and then the pellets were washed thrice using K-medium (0.032 M KCl, 0.051 M NaCl) (Williams and Dusenbery, 1990). The collected embryos were placed onto a new NGM plate, and incubated for 2 days (L4 stage) for soil toxicity tests.

### Short-term Exposure Test

We added 0.5 g of test soils into each well of a 24-well plate, together with 116 μL of K-medium (WHC_80%_-20 μL) (n=6). In order to prepare a food medium, we inoculated *E. coli* (strain OP50) into 50 mL of Luria-Bertani medium (LB broth, 2.5 g L^-1^) under sterile conditions, and incubated them overnight at 36±2°C (OD_600_: 0.8–1.0). We centrifuged 5 mL of cell suspension at 5,000 rpm for 20 min, and washed trice using K-medium. We resuspended the cell pellet using 1 mL of K-medium, and added 10 μL of cholesterol solution (cholesterol 5 mg mL^-1^ in >96% ethanol) (ISO, 2010). We added 20 μL of the food medium into each well of a 24-well plate, and ten age-synchronized nematodes were added to each well and maintained at 20 ± 2°C in the dark. We filled 10 mL of deionized water into the hollow space of the 24-well plate in order to prevent water loss during incubation periods, and sealed the plates with laboratory Parafilm. After 2 days, 1 mL of K-medium was added into a well, and suspended using a pipette. The supernatant was immediately (within 2 sec) removed, and the adult nematodes in the soil suspension were collected under the microscope. We repeated this process until no nematode was observed anymore (maximum 3 times), and recorded the survival rate (%) to initially exposed nematodes (n=6). In order to measure growth (length of the body) and brood size (number of embryos in the body), we randomly sampled 30 adults, and each body length was measured using an image analyzer (ImageJ, 1.52a, National Institutes of Health, USA). The data were expressed as a percentage (%) of the average value of the control group.

### Lifetime Exposure Test

The lifespans of *C. elegans* in the soil can be different from those in culture medium (Van Voorhies et al., 2005). We found that the survival rate of nematodes in the soil media decreased faster than in NGM (n=8) (Figure S2), and determined the lifetime exposure period to be 12 days (from embryo) since the survival rate at this time started to be reduced by half (about 50%). We used 2 day-old nematodes for toxicity tests, and the actual period of the lifetime exposure was 10 days. In order to perform the lifetime exposure test, we prepared five TWP-soil plates (24-well plates) equivalently to the short-term exposure test. These TWP-soil plates contain TWP-soil mixtures of each concentration, and are moistened by K-medium with a process equivalent to the short-term exposure test. The food medium was added into the first plate, and ten age-synchronized nematodes were added to each well. The second to fifth plates were first incubated without the food medium and nematodes. Since newly born juveniles in test soils become adults within 3 days, we isolated the initially exposed adults every 2 days to distinguish them from the grown adults in the soil. After 2 days (2^nd^ day after first exposure), the number of survivors in the first plate were recorded, and then re-exposed to the second plate after adding the food medium (n=6). This process was repeated every 2 days using third to fifth plates, so that the nematodes were exposed to TWP-soil mixtures that had experienced an increasing number of days of incubation (Figure S3). The survival rates (%) on the 4^th^ to 10^th^ day after first exposure were calculated.

### Pre-soil Incubation

In order to evaluate the pre-incubation time-dependent changes on TWP toxicity, we prepared two TWP-soil plates for short-term exposure tests. They were incubated without the food medium for two periods (30 and 75 days) at 20 ± 2°C in the dark. After soil pre-incubations, the plates were dried at room temperature in the sterile hood for 16 h, and 116 μL of deionized water was added into each well. Then short-term exposure tests were performed using each plate. For the lifetime exposure test, five TWP-soil plates were incubated for one period (75 days), and the plates were dried and moistened with deionized water. The lifetime exposure test was conducted with an equivalent process (see methods). The pH of each soil after each pre-incubation period ranged from 6.85 to 7.12 (Figure S4), which is in the optimal pH range (3.2 to 11.8) for nematode survival (Khanna et al., 1997).

### Water-extractable Metals in the TWP-soils

We prepared two TWP-soil plates for chemical analysis, and the plates were incubated without the food medium for 30 and 75 days at 20 ± 2°C in the dark, after which they were dried at room temperature in the sterile hood. We determined water-extractable metal concentrations in the TWP-soil mixtures as a proxy for bioavailable metal concentrations. The results of extractable (available) metal concentration highly depend on methodology (e.g. extraction solution) (Peijnenburg et al., 2007), but we here preferred a short extraction, as longer extraction times can lead to additional chemical leaching, thus confounding our result. In order to measure water-extractable metal concentrations in each soil, each soil sample was placed into 15 mL-test tubes, and 3 mL of K-medium was added into the tubes. We vortexed the tubes for 30 secs, and centrifuged at 5,000 rpm for 20 min. The supernatant was passed through a syringe filter (pore size 0.45 μm; D-76185, ROTILABO®, Carl Roth GmbH & Co., Karlsruhe, Germany), and stored at 4°C until analysis. The water-extractable metal (Cr, Cu, Ni, Pb, and Zn) concentrations in the solutions were measured by inductively coupled plasma optical emission spectrometer (ICP-OES, Perkin Elmer Optima 2100DV), and converted to soil concentration (mg kg^-1^).

### Statistical Analysis

Data were analyzed using the OriginPro software (OriginPro 8 SR2, Ver. 8.0891, OriginLab Corporation, MA, USA). One-way analysis for variance (ANOVA) and Turkey’s tests were conducted to determine the significance (*p* < 0.05) of multiple comparisons.

## RESULTS AND DISCUSSION

### Short-term Exposure Test

There was no effect on survival rate after the short-term exposure period (2 days, Figure 2A), but significant inhibition in growth started at 100 mg kg^-1^ of TWP concentration (Figure 2B). After 30 and 75 days of soil pre-incubation, the effects on growth were observed at lower concentrations (1 mg kg^-1^), and the growth inhibition rates in TWP treatment groups (1–10,000 mg kg^-1^) were 9–17% compared with the control (Figure 2B and S5A). Brood sizes significantly diminished at 1,000 to 10,000 mg kg^-1^ of TWP concentration (Figure 2C). After 30 and 75 days of soil pre-incubation, toxic effects occurred even at the lowest concentration (1 mg kg^-1^), and brood sizes were reduced 38–77% compared with the control. The ratios of non-pregnant individuals increased with soil pre-incubation time: 3–13% (no pre-incubation), 7–17% (30 days), and 20–33% (75 days) (Figure 2C and S5B).

**Figure 2.**
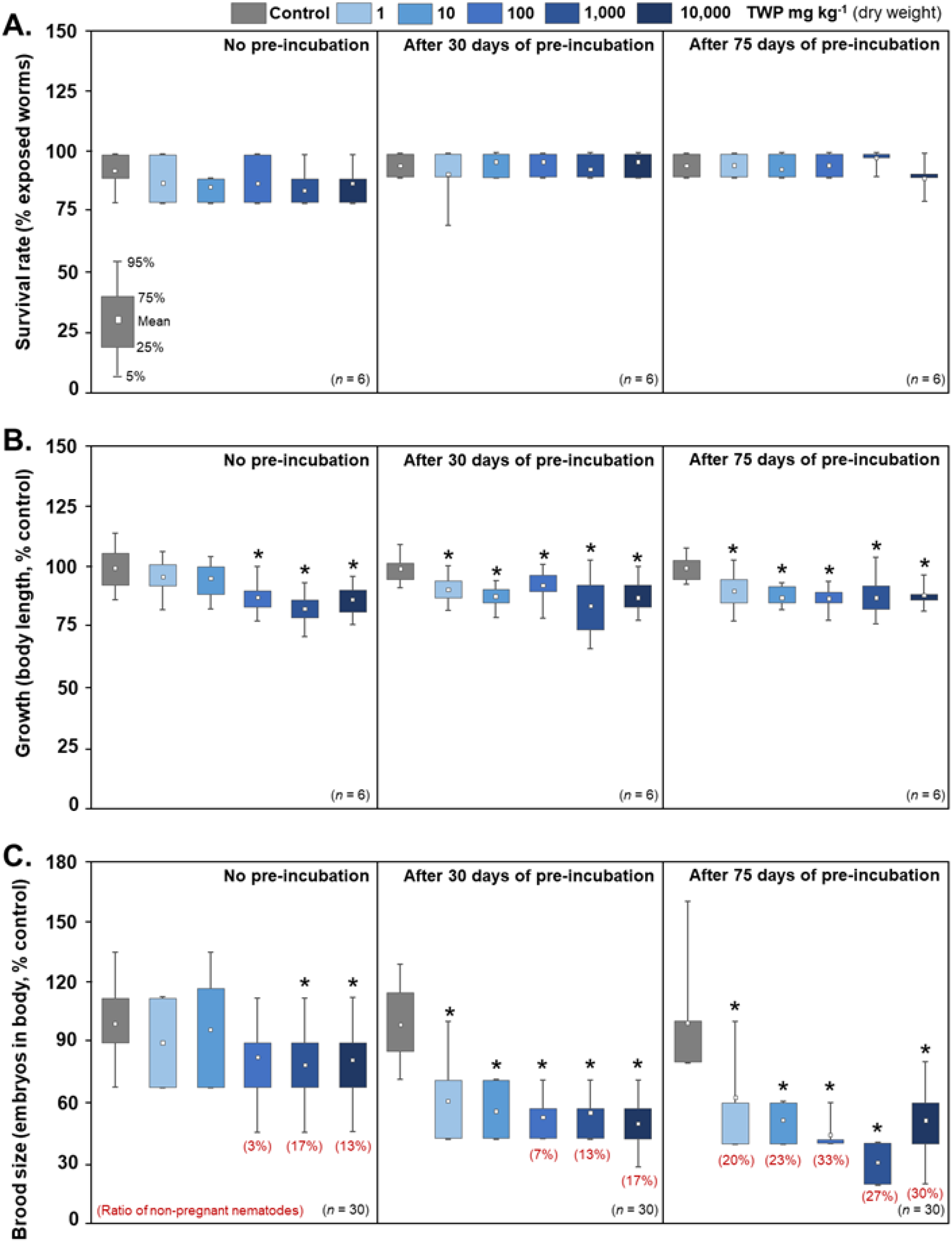
Toxic effects of TWPs on (A) survival, (B) growth (body of length), and (C) brood size (number of embryos in body) of *C. elegans* after short-term exposure period (2 days). The short-term exposure tests were performed after three different soil pre-incubation times (no pre-incubation, 30 days, and 75 days). The survival rate (%) was calculated to initially exposed nematodes, and the data of growth and brood size were normalized to each control group (% control). The asterisks (*) indicate significant (*p* < 0.05) differences compared to the control treatment group.

The ingestion of microplastics has been proposed as a main exposure route (Rehse et al., 2016), but we avoided nematodes feeding on microplastics since the size range of TWPs used here (34–265 μm) was much larger than the edible size (≤3.4 μm) (Fueser et al., 2019; Mueller et al., 2020). Significant effects of TWPs on growth and brood size were observed at 100 and 1,000 mg kg^-1^, which are lower than the effects of other microplastic fragments (e.g. high-density polyethylene, polyethylene terephthalate, polypropylene, and polystyrene) on *C. elegans* in the soil (Kim et al., 2020). Other soil organisms have also been shown to be negatively affected by TWPs: The growth of the earthworm (*Eisenia fetida*) was reduced in soil mixtures containing virgin crumb rubber (50:50) after a 33 day exposure period (Pochron et al., 2017), whereas no effect was observed in black worms (*Lumbriculus variegatus*) at <10% of tire wear particles (based on dry sediment) (Redondo-Hasselerharm et al., 2018). These data cannot be directly compared with our results due to the different experimental conditions (e.g., tire composition or virgin versus aged material), but our TWP sample seems to have high toxicity on the nematodes.

After 30 and 75 days of soil pre-incubation, the adverse effects on nematodes occurred at the lowest concentration (1 mg kg^-1^), indicating that soil pre-incubation increased the toxicity of the TWPs. This level is much lower than environmentally relevant concentrations of tire wear particles reported from roadside soils (400 to 158,000 mg kg^-1^) (Kocher et al., 2008). This result is in agreement with our previous study reporting high toxicity on plants at very low TWP concentrations (same material as used in this study, 10 mg kg^-1^) (Leifheit et al., 2021). We here assumed that the increase of toxicity after soil pre-incubation may be strongly linked with chemicals leaching out of the TWPs. Artificially or naturally aged microplastics generally induce higher toxicity than pristine ones since the fragmentation and chain-scission of microplastics by aging result in increased release of additives (Liu et al., 2021). Recent studies used microplastics after aging or extraction treatment, and found that the treated microplastic particles lose their toxicity due to the loss of extractable chemicals (Kim et al., 2020; Pflugmacher et al., 2021). In previous papers regarding tire toxicity, the metal concentration (Zn) in water containing tire wear particles increased with time (5 to 45 days), and it was bioavailable to organisms (Camponelli et al., 2009; Turner and Rice, 2010). The threshold value (1 mg kg^-1^) of our study is a rarely found level in nematode toxicity tests. For instance, 50% effective concentrations (EC50) of heavy metals (As, Cd, Cu, Ni, and Zn) on reproduction are calculated in the range of 303 to 744 mg kg^-1^ (Kim et al., 2018). We speculate that the nematodes may have been influenced by synergistic effects of various chemicals (metals, PAHs, and antioxidants) released from TWPs during soil pre-incubation and hence chose this low concentration. Previous studies reported that synergistic effects can occur in 25 mixtures of five biocides and five heavy metals (Xu et al., 2011), and the pairwise or triple exposure of various metals can display synergistic killing effects on *C. elegans* (Chu et al., 2002).

### Lifetime Exposure Test

The significant reduction in the lifespan of *C. elegans* started on the 8^th^ day after first exposure, and the survival times of TWP treatment groups on the 10^th^ day were reduced 27–45% compared with the control (Figure 3A). These effects occurred at all TWP concentrations including the lowest one (1 mg kg^-1^), which showed no toxicity in short-term exposure tests targeting growth (100 mg kg^-1^) and brood size (1,000 mg kg^-1^) (Figure 2B and 2C). After 75 days of soil pre-incubation, a significant reduction occurred on the 6^th^ day, and the survival rates of TWP treatment groups were 17–50% on the 10^th^ day (Figure 3B).

**Figure 3.**
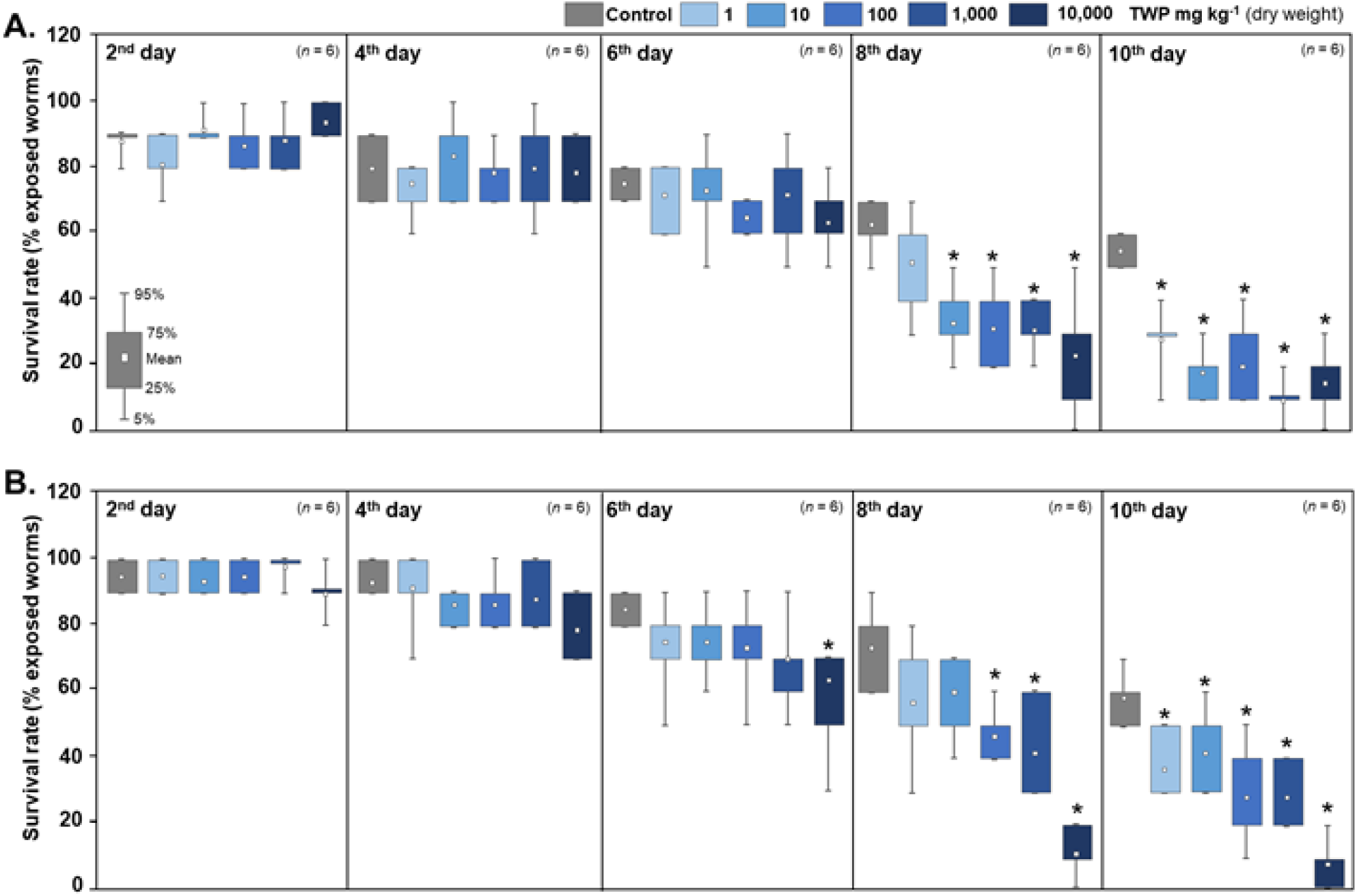
Survival rates of *C. elegans* after the lifetime exposure tests. The lifetime exposure tests were performed after two different soil pre-incubation times: (A) no pre-incubation and (B) 75 days. The survival rate (%) was calculated to initially exposed nematodes at every 2 days until 10^th^ day after first exposure. The asterisks (*) indicate significant (*p* < 0.05) differences compared to the control treatment group.

The exposure period of the toxicity test is generally standardized since it can reduce variation in toxicity sensitivity resulting from different exposure times, and guarantee better data fit for ecological risk assessment. However, the standardized toxicity test targets a certain period of the test species’ lifespan. For instance, the chronic exposure period for nematode *C. elegans* is generally 4 days, even though their lifespan is 18–20 days at 20°C in the culture media (Amrit et al., 2014; ISO, 2020). In order to reflect more environmentally relevant conditions, previous papers have used the lifespan test (or continuous exposure test), and found that exposure time can highly influence the extent of toxic effects (Gonçalves et al., 2017; Peters and Granek, 2016). In microplastic research, this approach can be an appropriate metric to reflect the spectrum of changes on microplastic toxicity over time (Shang et al., 2020). The toxic effects of microplastics will be gradually higher with time due to time-delayed release of chemical additives (Rillig et al., 2021), while target organisms will be eventually older and potentially more susceptible. A recent study conducted a lifetime exposure (103 days) test using a common freshwater invertebrate (*Daphnia magna*), and found that sub-lethal concentrations of polystyrene nanoparticles were two orders of magnitude lower than the concentration in the short-term exposure test (1 day) (Kelpsiene et al., 2020). The lifespan assay using *C. elegans* was also performed in many previous studies, and microplastic exposure has been shown to potentially shorten their life expectancy (Lei et al., 2018; Shang et al., 2020). This assay has been widely conducted in culture medium (agar or liquid) (Amrit et al., 2014), but not in soil media, and our study is the first to estimate toxicity during lifetime of *C. elegans* in the soil system. In previous papers regarding tire toxicity, there was no available data from lifetime tests, but only from chronic tests using aquatic organisms. Chronic exposure (21 days) of tire wear particles (0.58 g L^-1^) can lead to high mortality (92.5%) of freshwater invertebrates while the 50% lethal concentration was 1 g L^-1^ after acute exposure (2 days) (Khan et al., 2019), but another study observed no effects on *Hyalella azteca* (amphipod) and *Ceriodaphnia dubia* (water flea) in tire wear particle-spiked sediment (10,000 mg kg^-1^) and its elutriates (Panko et al., 2013) after chronic exposure test. We here found a reduction of survival time at even the lowest TWP concentration (1 mg kg^-1^), which is 100- to 1,000-folds lower than the effective concentrations on growth and brood size. After 75 days of soil pre-incubation, the nematodes in the TWP treatment groups started to die earlier than in the no pre-incubation treatment, and this may be related with the different initial concentration of chemicals released from TWP after soil pre-incubation.

### Water-extractable Metals in the TWP-soils

The water-extractable (and thus likely bioavailable) concentrations of Cr, Cu, Ni, Pb, and Zn in the TWP-soils did not vary among treatments (TWP concentrations and pre-incubation times) (Table S1). We expected that water-extractable metal concentrations in TWP-soils would increase with time, but there was no correlation with nominal tire concentrations (1–10,000 mg kg^-1^) or pre-incubation times (no pre-incubation, 30 days, and 75 days). This implies that either the added TWPs did not contain sufficient quantities of metals to cause a measurable change, or our test period was not long enough or the soil pH was too high to release metals from TWPs. We conclude that the metals incorporated in TWPs are likely not the main reason of toxicity in this study, but other chemical additives such as PAHs or antioxidants may be linked with high toxicity of TWPs.

We found 19 previous publications focusing on the effects of tire materials (Table S2). These papers have claimed that chemical leaching from tire materials is an important mediator of toxicity, but only 10 papers performed chemical analysis to screen chemical compositions in tire and its leachates. Five among these 10 papers targeted only Zn as a target chemical since it has been pointed out as a main reason for toxicity of tire leachates (Nelson et al., 1994), and Zn concentration showed a large variation depending on tire brands and pH of test media (Gualtieri et al., 2005; Wik et al., 2006). Another five papers analyzed PAHs, metals, organic compounds, and major nutrients in sediment- or soil-tire mixtures. However, several studies directly applied acid-based extraction procedures (e.g., HNO_3_ treatment) to quantify total chemical concentration in test media (Camponelli et al., 2009; Pochron et al., 2017; 2018; Redondo-Hasselerharm et al., 2018). Although they observed high levels of target chemicals, their toxic potential is likely overestimated since strong acid treatment induces particle damage and chemical leaching (Vandermeersch et al., 2015). Various metals in the leachates of tire-sediment were detected by extraction at 44°C, and showed significant toxic effects on *Daphnia* species, but not at 21°C (Marwood et al., 2011). A recent study investigated total and CaCl_2_-extractable (bioavailable) metal contents in tire-sediment mixture (Redondo-Hasselerharm et al., 2018). Total Zn concentration (extracted by an acid-based method) showed a linear correlation with nominal tire concentration in the sediment, but CaCl_2_-extractable Zn was not detected in the same sediment. The authors of this study claimed that tire material does not negatively affect freshwater benthic invertebrates. Based on this evidence, the bioavailability of the released tire-chemicals in the solid media such as soil or sediment may be highly important in tire particle toxicity since metal adsorption onto soil particles is a major process responsible for bioaccumulation (Peijnenburg et al., 2007). In addition, mild extractions are probably more appropriate for estimating bioavailable chemicals in the soil solution, which will be predominantly responsible for toxic effects. The contradictory findings of these 19 studies can be attributed to the use of different methods to determine chemical components, different soil with differing pH, and the use of different target organisms.

Our ecotoxicological data show that TWPs can be very harmful in the soil environment at low concentrations, and it is vital to pay more attention to this issue in future toxicity tests. Although our study was performed on a small scale, we clearly observed a time-delayed toxicity in TWP-soil mixtures, and this toxic effect can occur at much lower concentrations than the environmentally relevant concentration for this pollutant. Toxic effects of microplastics will be gradually higher with exposure time, and susceptibility of target organisms likely changes with developmental stage; taken together, this implies that microplastic toxicity tests need to embrace a variety case of developmental and perhaps also transgenerational levels. We suggest that exposure time should be considered as one of the main factors in evaluating microplastic toxicity, and long-term studies should be conducted with environmentally relevant concentrations. Future tests should focus on finding correlations between a key explanatory chemicals or chemical mixtures and tire toxicity in the soil.

## ASSOCIATED CONTENT

### AUTHOR INFORMATION

#### Supporting Information

The Supporting information is available free of charge on the ACS Publications website.

■ Methods: FTIR spectra of each target MP (Figure S1), Lifespan of *C. elegans* in soil and NGM (Figure S2), A diagram depicting the lifetime exposure test (Figure S3), Soil pH in each TWP-soil mixture (Figure S4)
■ Result and Discussion: Pictures of *C. elegans* after short-term exposure tests (Figure S5), Water-extractable metal concentrations in TWP-soil mixtures (Table S1), List of previous studies reporting tire toxicity (Table S2).

#### Notes

The authors declare no competing financial interest.

## ACKNOWLEDGEMENTS

MCR acknowledges support from an ERC Advanced Grant (grant no. 694368). SM acknowledges support by the German Federal Ministry of Education and research (BMBF) within the collaborative Project “Bridging in Biodiversity Science (BIBS-phase 2)” (funding number 01LC1501). EFL acknowledges funding from the Deutsche Forschungsgemeinschaft (LE 3859/1-1).

## Notes

### Competing Interest Statement

The authors have declared no competing interest.

