## Supplementary Information for "Time-dependent toxicity of tire particles on soil nematodes"

11 pages

5 figures

2 tables

Reference list

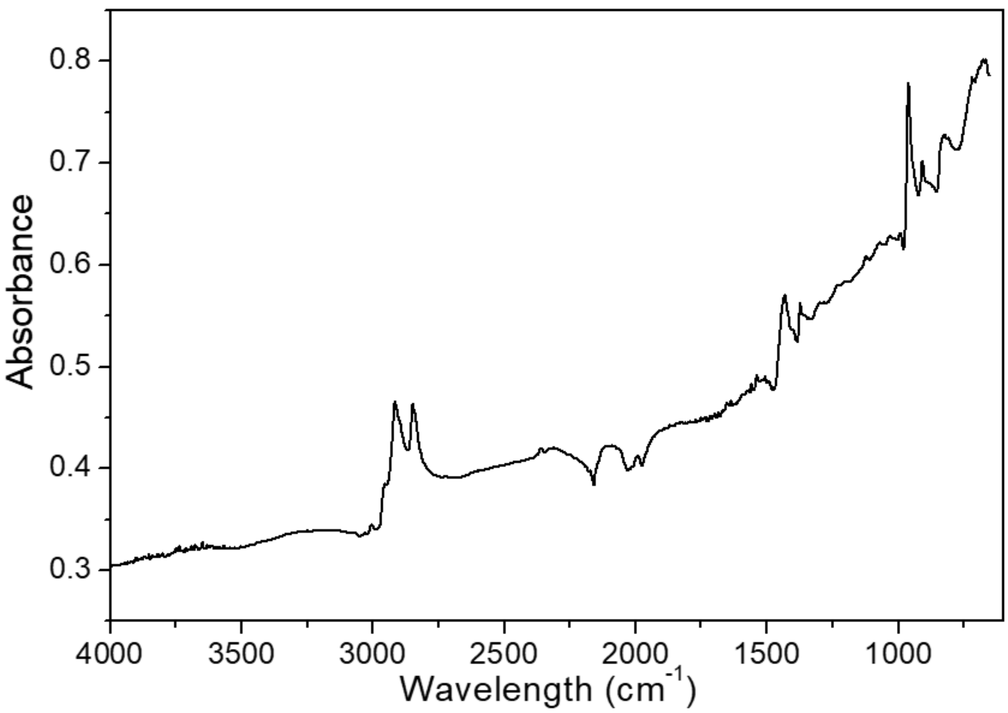


**Figure S1.** FTIR spectra, ATR (attenuated total reflection) mode, of TWP tested in this paper

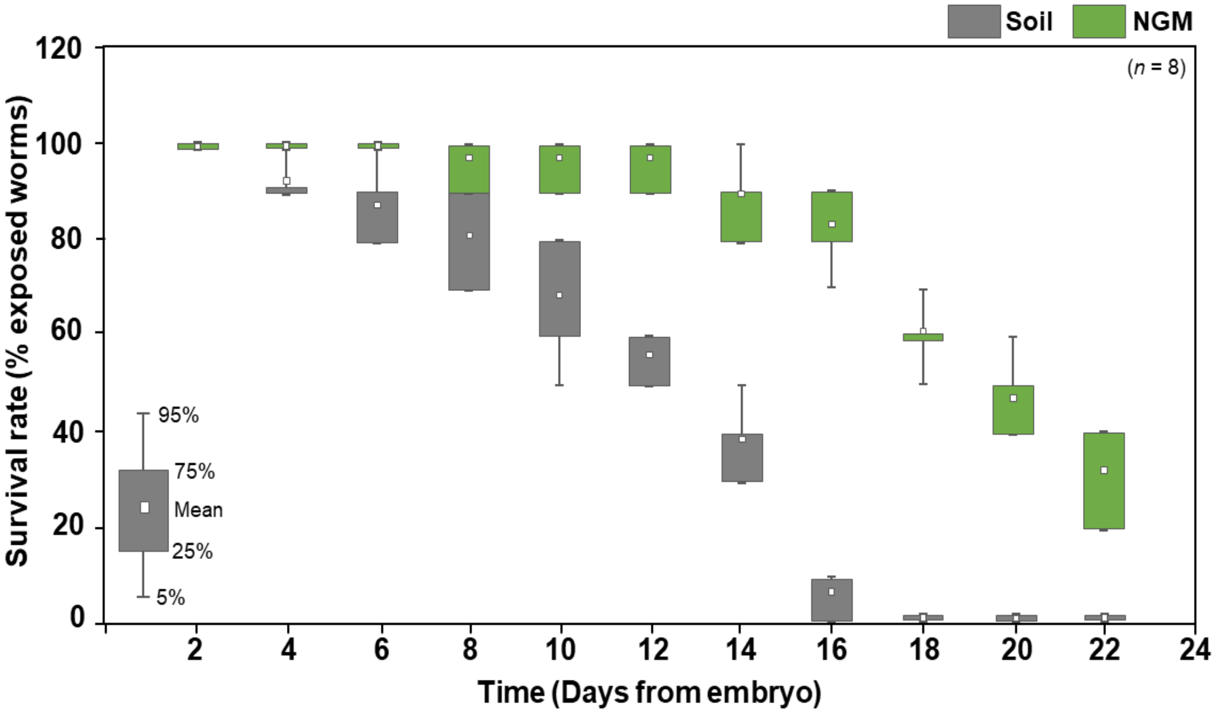

**Figure S2.** The lifespan of *C. elegans* in soil and NGM.

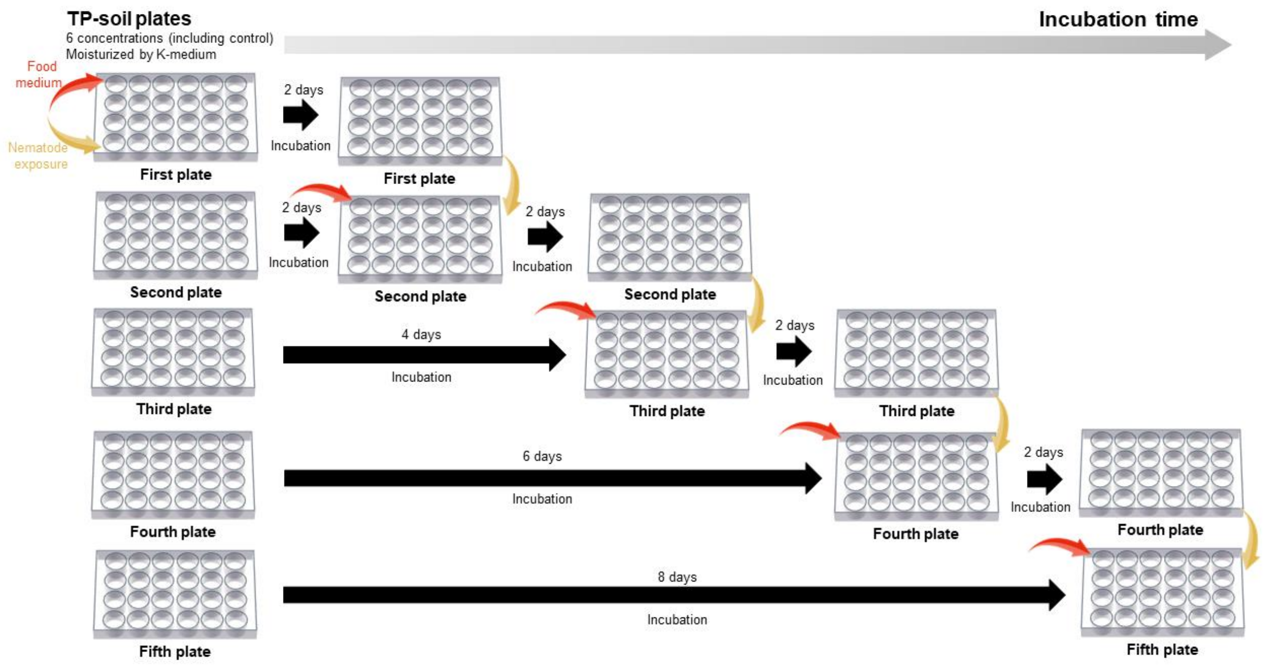


**Figure S3.** The diagram depicting the lifetime exposure test in this study.

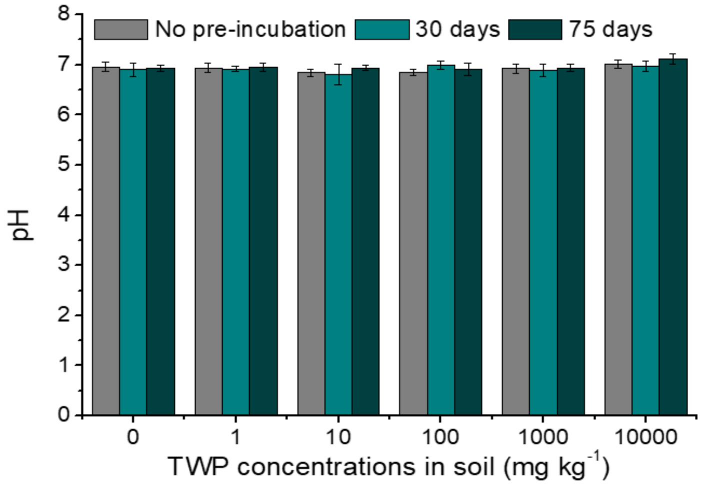


**Figure S4.** Soil pH in each tested soil (six concentrations and three soil pre-incubation times) in this study.

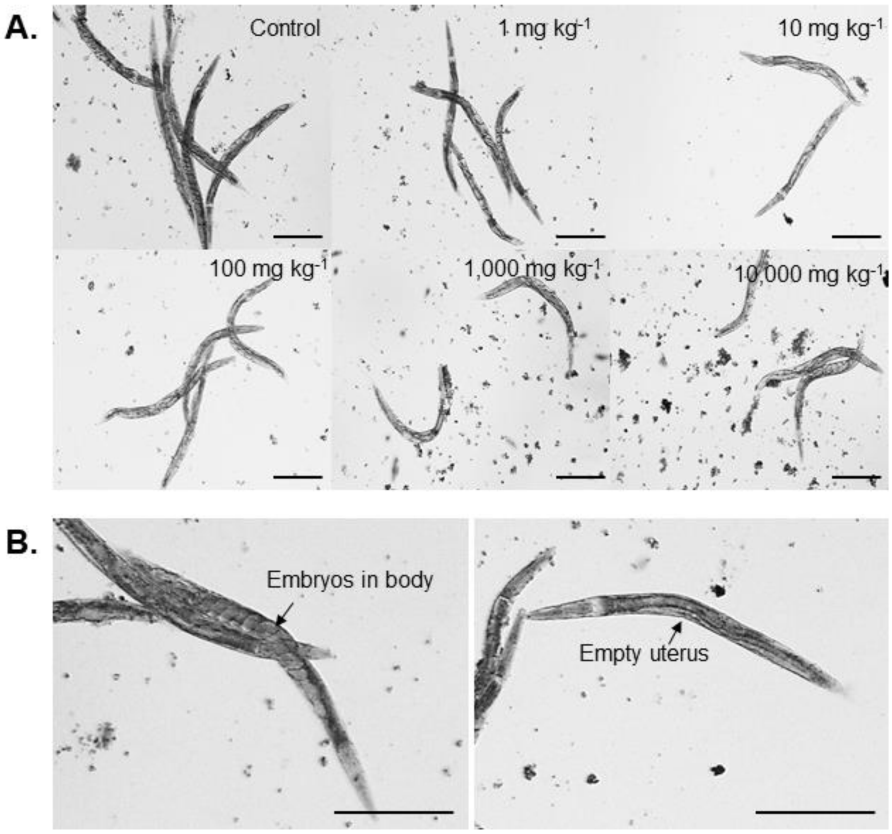


**Figure S5.** The collected *C. elegans* from TWP-soil mixtures after short-term exposure test (75 days of soil pre-incubation): (A) different body length of *C. elegans* from TWP-soil mixtures of six different concentrations, (B) pregnant (*left*) and non-pregnant (*right*) nematodes (scale bar: 250 μm)

**Table S1.** The water-extractable metal (Cr, Pb, Zn, Ni, Cu) concentration (mg $kg^{-1}$) in each soil (six concentrations and three soil pre-incubation times).

| | **Control** | **1 mg $kg^{-1}$** | **10 mg $kg^{-1}$** | **100 mg $kg^{-1}$** | **1,000 mg $kg^{-1}$** | **10,000 mg $kg^{-1}$** |
|---|---|---|---|---|---|---|
| **Cr (mg $kg^{-1}$)** | | | | | | |
| **No pre-incubation** | | | | | | |
| 1 | 0.005 | n.d. | 0.002 | 0.002 | n.d. | n.d. |
| 2 | 0.004 | n.d. | n.d. | 0.007 | n.d. | n.d. |
| 3 | 0.000 | n.d. | n.d. | 0.003 | n.d. | n.d. |
| 4 | 0.001 | n.d. | 0.000 | 0.001 | n.d. | n.d. |
| *Average* | *0.002* | *n.d.* | *0.001* | *0.003* | *n.d.* | *n.d.* |
| *Stdev* | *0.002* | *n.d.* | *0.001* | *0.002* | *n.d.* | *n.d.* |
| **30 days** | | | | | | |
| 1 | n.d. | n.d. | n.d. | n.d. | n.d. | n.d. |
| 2 | n.d. | 0.002 | n.d. | n.d. | n.d. | n.d. |
| 3 | n.d. | n.d. | n.d. | n.d. | n.d. | n.d. |
| 4 | n.d. | n.d. | n.d. | n.d. | n.d. | n.d. |
| *Average* | *n.d.* | *0.002* | *n.d.* | *n.d.* | *n.d.* | *n.d.* |
| *Stdev* | *n.d.* | *-* | *n.d.* | *n.d.* | *n.d.* | *n.d.* |
| **75 days** | | | | | | |
| 1 | 0.003 | n.d. | n.d. | n.d. | n.d. | n.d. |
| 2 | 0.005 | 0.002 | n.d. | n.d. | n.d. | n.d. |
| 3 | n.d. | n.d. | n.d. | n.d. | n.d. | n.d. |
| 4 | n.d. | n.d. | n.d. | n.d. | n.d. | n.d. |
| *Average* | *0.004* | *0.002* | *n.d.* | *n.d.* | *n.d.* | *n.d.* |
| *Stdev* | *0.002* | *-* | *n.d.* | *n.d.* | *n.d.* | *n.d.* |
| **Pb (mg $kg^{-1}$)** | | | | | | |
| **No pre-incubation** | | | | | | |
| 1 | n.d. | 0.001 | 0.011 | 0.020 | 0.015 | 0.002 |
| 2 | 0.015 | 0.000 | 0.018 | 0.018 | n.d. | n.d. |
| 3 | 0.013 | 0.014 | n.d. | 0.046 | n.d. | n.d. |
| 4 | n.d. | 0.026 | 0.012 | 0.018 | n.d. | 0.011 |
| *Average* | *0.014* | *0.010* | *0.013* | *0.025* | *0.015* | *0.007* |
| *Stdev* | *0.001* | *0.010* | *0.003* | *0.012* | *0.000* | *0.004* |
| **30 days** | | | | | | |
| 1 | n.d. | n.d. | 0.015 | 0.005 | n.d. | 0.001 |
| 2 | 0.019 | 0.009 | n.d. | 0.020 | 0.003 | 0.025 |
| 3 | n.d. | 0.008 | 0.009 | 0.004 | n.d. | 0.003 |
| 4 | n.d. | 0.002 | n.d. | n.d. | 0.008 | 0.004 |
| *Average* | *0.019* | *0.006* | *0.012* | *0.010* | *0.005* | *0.008* |
| *Stdev* | *0.000* | *0.003* | *0.003* | *0.007* | *0.003* | *0.010* |
| **75 days** | | | | | | |
| 1 | 0.006 | n.d. | 0.016 | 0.034 | 0.029 | 0.030 |
| 2 | 0.011 | n.d. | 0.012 | 0.029 | 0.037 | 0.032 |
| 3 | n.d. | 0.011 | 0.018 | 0.030 | 0.023 | 0.026 |
| 4 | n.d. | n.d. | 0.035 | 0.036 | 0.032 | 0.025 |
| *Average* | *0.008* | *0.011* | *0.020* | *0.032* | *0.030* | *0.028* |
| *Stdev* | *0.002* | *0.000* | *0.009* | *0.003* | *0.005* | *0.003* |
| **Zn (mg $kg^{-1}$)** | | | | | | |
| **No pre-incubation** | | | | | | |
| 1 | 0.027 | 0.016 | 0.015 | 0.018 | 0.059 | 0.094 |
| 2 | 0.035 | 0.019 | 0.019 | 0.018 | 0.029 | 0.119 |
| 3 | 0.019 | 0.017 | 0.015 | 0.050 | 0.022 | 0.125 |
| 4 | 0.030 | 0.025 | 0.019 | 0.027 | 0.022 | 0.106 |
| *Average* | *0.028* | *0.019* | *0.017* | *0.028* | *0.033* | *0.111* |
| *Stdev* | *0.006* | *0.004* | *0.002* | *0.013* | *0.015* | *0.012* |
| **30 days** | | | | | | |
| 1 | 0.010 | 0.013 | 0.012 | 0.012 | 0.020 | 0.111 |
| 2 | 0.012 | 0.016 | 0.016 | 0.010 | 0.014 | 0.188 |
| 3 | 0.014 | 0.016 | 0.020 | 0.012 | 0.015 | 0.177 |
| 4 | 0.019 | 0.016 | 0.013 | 0.011 | 0.019 | 0.154 |
| *Average* | *0.014* | *0.015* | *0.015* | *0.011* | *0.017* | *0.157* |
| *Stdev* | *0.003* | *0.001* | *0.003* | *0.001* | *0.003* | *0.030* |
| **75 days** | | | | | | |

| | | | | | | |
|---|---|---|---|---|---|---|
| 1 | 0.018 | 0.010 | n.d. | n.d. | n.d. | n.d. |
| 2 | 0.017 | 0.019 | n.d. | n.d. | n.d. | n.d. |
| 3 | 0.043 | n.d. | n.d. | n.d. | n.d. | n.d. |
| 4 | 0.029 | n.d. | n.d. | n.d. | n.d. | n.d. |
| *Average* | *0.027* | *0.015* | *n.d.* | *n.d.* | *n.d.* | *n.d.* |
| *Stdev* | *0.011* | *0.004* | *n.d.* | *n.d.* | *n.d.* | *n.d.* |
| **Ni (mg kg$^{-1}$)** | | | | | | |
| **No pre-incubation** | | | | | | |
| 1 | 0.013 | 0.029 | 0.004 | 0.004 | 0.024 | 0.003 |
| 2 | n.d. | 0.002 | 0.004 | n.d. | 0.005 | 0.004 |
| 3 | 0.002 | n.d. | 0.001 | 0.001 | n.d. | 0.040 |
| 4 | 0.012 | n.d. | 0.006 | 0.004 | n.d. | 0.009 |
| *Average* | *0.009* | *0.016* | *0.003* | *0.003* | *0.015* | *0.014* |
| *Stdev* | *0.005* | *0.013* | *0.002* | *0.002* | *0.009* | *0.015* |
| **30 days** | | | | | | |
| 1 | 0.002 | 0.001 | 0.006 | 0.006 | 0.002 | 0.003 |
| 2 | n.d. | 0.001 | 0.005 | 0.001 | 0.003 | 0.004 |
| 3 | 0.002 | n.d. | n.d. | 0.001 | 0.007 | 0.001 |
| 4 | 0.001 | 0.005 | 0.005 | 0.002 | 0.000 | 0.003 |
| *Average* | *0.002* | *0.003* | *0.005* | *0.002* | *0.003* | *0.003* |
| *Stdev* | *0.001* | *0.002* | *0.001* | *0.002* | *0.002* | *0.001* |
| **75 days** | | | | | | |
| 1 | 0.005 | 0.005 | n.d. | n.d. | n.d. | n.d. |
| 2 | 0.004 | 0.009 | n.d. | n.d. | n.d. | n.d. |
| 3 | 0.006 | n.d. | n.d. | n.d. | n.d. | n.d. |
| 4 | 0.006 | n.d. | n.d. | n.d. | n.d. | n.d. |
| *Average* | *0.005* | *0.007* | *n.d.* | *n.d.* | *n.d.* | *n.d.* |
| *Stdev* | *0.001* | *0.002* | *n.d.* | *n.d.* | *n.d.* | *n.d.* |
| **Cu (mg kg$^{-1}$)** | | | | | | |
| **No pre-incubation** | | | | | | |
| 1 | 0.106 | 0.072 | 0.065 | 0.049 | 0.068 | 0.058 |
| 2 | 0.118 | 0.056 | 0.048 | 0.041 | 0.178 | 0.063 |
| 3 | 0.065 | 0.048 | 0.059 | 0.047 | 0.044 | 0.089 |
| 4 | 0.121 | 0.079 | 0.101 | 0.044 | 0.041 | 0.076 |
| *Average* | *0.103* | *0.064* | *0.068* | *0.045* | *0.083* | *0.071* |
| *Stdev* | *0.022* | *0.012* | *0.020* | *0.003* | *0.056* | *0.012* |
| **30 days** | | | | | | |
| 1 | 0.029 | 0.037 | 0.022 | 0.241 | 0.018 | 0.013 |
| 2 | 0.027 | 0.031 | 0.022 | 0.043 | 0.018 | 0.016 |
| 3 | 0.023 | 0.029 | 0.015 | 0.018 | 0.020 | 0.014 |
| 4 | 0.030 | 0.026 | 0.022 | 0.019 | 0.019 | 0.026 |
| *Average* | *0.027* | *0.031* | *0.020* | *0.080* | *0.019* | *0.017* |
| *Stdev* | *0.003* | *0.004* | *0.003* | *0.093* | *0.001* | *0.005* |
| **75 days** | | | | | | |
| 1 | 0.029 | 0.019 | 0.046 | 0.033 | 0.032 | 0.033 |
| 2 | 0.038 | 0.035 | 0.046 | 0.032 | 0.032 | 0.033 |
| 3 | 0.078 | 0.045 | 0.032 | 0.032 | 0.033 | 0.032 |
| 4 | 0.074 | n.d. | 0.032 | 0.033 | 0.033 | 0.033 |
| *Average* | *0.054* | *0.033* | *0.039* | *0.032* | *0.032* | *0.033* |
| *Stdev* | *0.022* | *0.011* | *0.007* | *0.000* | *0.000* | *0.001* |

**Table S2.** List of previous studies reporting tire toxicity.

| No. | Year | Author | Type | Tire characterization | | Media | Target species | Effects | Chemical analysis | |
|---|---|---|---|---|---|---|---|---|---|---|
| 1 | 2005a | Gualtieri et al. | Tire-leachate | Tire scraps | 1–7 µm | Water | *Xenopus laevis* | Teratogenic effects | No | - |
| 2 | 2007 | Mantecca et al. | Tire-leachate | Tire scraps | - | FETAX solution | *Xenopus laevis* | Embryotoxicity | No | - |
| 3 | 1998 | Hartwell et al. | Tire | Tire scraps | - | Synthetic seawater | *Cyprinodon variegatus, Palaemonetes pugio* | Correlation between toxicity and salinity | No | - |
| 4 | 2000 | Hartwell et al. | Tire | Tire chips | 1 cm3 | Synthetic seawater | *Allivibrio fischeri* | Correlation between toxicity and salinity | No | - |
| 5 | 2003 | Stephensen et al. | Tire | Whole tire | - | Freshwater | *Oncorhynchus mykiss* | Effects on Ethoxyresorufin-O-deethylase activity | No | - |
| 6 | 2005 | Wik and Dave | Tire | Grinded tire | - | Water | *Daphnia magna* | Immobility | No | - |
| 7 | 2013 | Panko et al. | Tire-leachate | Road simulator particle | <150 µm | Freshwater sediment | *Ceriodaphnia dubia, Pimephales promelas, Chironomus dilutus, Hyalella azteca* | No effects | No | - |
| 8 | 2019 | Khan et al. | Tire | Grinded tire | <500 µm | Water | *Hyallela azteca* | Effects on mortality, growth, reproduction | No | - |
| 9 | 2021 | Leifheit et al. | Tire | Grinded tire | 34-265 µm | Soil | *Allium porrum* | Effects on plant biomass | No | - |
| 10 | 2005b | Gualtieri et al. | Tire-leachate | Road simulator particle | 10–80 µm | Water | *Raphidocelis subcapitata, Daphnia magna, Xenopus laevis* | Teratogenic effects, sub-lethal effects | Yes | Zn |
| 11 | 2006 | Wik and Dave | Tire | Abraded tire | - | Water | *Daphnia magna* | Immobility | Yes | Zn |
| 12 | 2009 | Wik et al. | Tire-leachate | Abraded tire | - | MilliQ water | *Daphnia magna, Ceriodaphnia dubia, Pseudokirchnerella subcapitata, Danio rerio* | Immobility on water flea, no effects on fish | Yes | Zn |
| 13 | 2010 | Turner and Rice | Tire-leachate | Abraded tire | <500 µm | Filtered sea water | *Ulva lactuca* | Effects on chlorophyll | Yes | Zn |
| 14 | 2017 | Villena et al. | Tire-leachate | Ground tire material | - | deionized water | *Aedes albopictus, Aedes triseriatus* | Effects of survival | Yes | Zn |
| 15 | 2009 | Camponelli et al. | Tire | Tire scraps | <590 µm | Sediment | *Rana sylvatica* | Significant difference in hatching success | Yes | Metals (Cr, Ni, Cu, Zn, As, Se, Cd, Pb) in sediment, water (extracted by $HNO_3$ at 150°C) |
| 16 | 2011 | Marwood et al. | Tire-leachate | Road simulator particle | <150 µm | Freshwater sediment | *Pseudokirchnerella subcapitata, Daphnia magna, Pimephales promelas* | Toxicity at high temp. leachates (44 °C) | Yes | Metals, PAHs, organic compounds present in tires-leachates |
| 17 | 2017 | Pochron et al. | Tire | Crumb rubber | - | Clean topsoil | *Eisenia fetida*, soil respiration | Effects on body weight | Yes | Major nutrients (Morgan extraction procedure) |
| 18 | 2018 | Pochron et al. | Tire | Crumb rubber | - | Clean topsoil | *Eisenia fetida* | No effects | Yes | Major nutrients (Morgan extraction procedure) |
| 19 | 2018 | Redondo-Hasselerharm et al. | Tire | Grinded tire | <500 µm | Freshwater sediments | *Gammarus pulex Asellus aquaticus, Tubifex spp, Lumbriculus variegatu* | No effects | Yes | Metals, PAHs in sediment (extractable, total) |
